# Dental pulp metagenomics reveals ancient human Herpesviridae

**DOI:** 10.1101/2022.04.06.487315

**Authors:** Oumarou Hama Hamadou, Zandotti Christine, Loukil Ahmed, Gazin Céline, Jonvel Richard, Georges-Zimmermann Patrice, Decker Michael, Fournier Pierre Edouard, Barbieri Rémi, Aboudharam Gérard, Drancourt Michel

## Abstract

Following a recent report on ancient human Herpesviridae (HHV), we adapted a standard protocol used for the high-throughput sequencing of modern DNA to recover ancient DNA (aDNA) from dental pulp specimens collected at two historical burial sites. Investigation of 36 soldiers buried in Sevastopol, Crimea, during the 1853-1856 Crimean War yielded genomic sequences covering 8.5% of the herpes simplex virus (HSV)-1 genome, 15.7% of the HSV-2 genome, 1.9% of the Epstein–Barr virus (EBV) genome and 3.9% of the cytomegalovirus genome. Further investigation of 16 civilians buried in the 17^th^ century in Amiens, France, yielded genomic sequences covering 9.8% of the HHV-1 genome and 2.2% of the HHV-6A genome; all data obtained with parallel negative controls was appropriately negative. Specific qPCRs confirmed the presence of four HSV-1, two HSV-2 and one EBV sequences in Sevastopol and two HSV-1 and one HHV-6 sequences in Amiens. HHV-6 was further detected by fluorescent *in situ* hybridization. These results demonstrated the usefulness of modern metagenomic protocols for high-throughput sequencing of aDNA in dental pulp.

---

A recent metagenomic-based study extended the antiquity of Herpesvirus in human populations after recovery of one partial and three complete genomes of human herpes simplex virus 1 (HSV-1) from European individuals dating from the 3rd-17^th^ centuries (Guellil *et al*. 2022). Such unprecedented results were obtained by the NextSeq500 sequencing system using extensive laboratory work adapted to ancient DNA (aDNA), which is known to be physically and chemically degraded (Keyser-Tracqui and Ludes 2004). Here, building on these recent data, we adapted a high-throughput sequencing protocol for modern DNA for screening for Herpesviridae aDNA in human samples collected from two archaeological sites. A war cemetery located in the Sevastopol suburbs, Crimea (global positioning system (GPS) coordinates = 44° 34 30″ N, 33° 26 23″ E), containing remains of likely French soldiers from the 1853-1856 Crimea War (Badem C., 2010) yielded 112 teeth from 36 individuals, consisting of 1-5 teeth per individual and 12 teeth not linked to any individual. Furthermore, a mass grave discovered within Hôtel-Dieu Hospital cemetery, Amiens, France (GPS = 49° 53 39″ N, 2° 17 45″ E), yielded 60 teeth from 16 individuals radiocarbon dated to the 17^th^ century, consisting of 1-5 teeth per individual. Both archaeological sites were investigated in agreement with regulations enforced at the time of investigations in Crimea and France. Dental pulp aDNA extracted from each tooth as previously described (Drancourt *et al*., 1998) was sequenced using the standard protocol for modern DNA and a NovaSeq sequencer (Illumina Inc., San Diego, CA, USA) (Appendix 1). All reads were mapped to reference genomes (HSV-1: NC_001806.2, HSV-2: NC_001798.2, varicella-zoster virus (VZV): NC_001348.1, Epstein–Barr virus (EBV): NC_007605.1, cytomegalovirus (CMV): NC_006273.2, HHV-6A: NC_001664.4, HHV-6B: NC_000898.1, HHV-7: NC_001716.2, HHV-8: NC_009333.1) using CLC Genomics Workbench software, version 7.5 (Qiagen). Metagenomic-based detection of Herpesviridae aDNA was tentatively confirmed by quantitative real-time polymerase chain reaction (qPCR) designed to amplify the internal control albumin gene and phage T4 and each metagenomic-detected HHV (Appendix 2).

No Herpesviridae DNA was detected in the negative control, which yielded only *Homo sapiens* sequences, but mapping Sebastopol reads detected 8.51% of the HSV-1 consensus genome (percentage of identical matches: 87.5-100%, depending on the alignment) in four individuals confirmed by qPCR; 15.71% of the HSV-2 consensus genome (percentage of identical matches: 85.04-100%) in two individuals confirmed by qPCR; 1.91% of the EBV consensus genome (percentage of identical matches: 94.11-97.77%) in one qPCR-positive individual; and 3.86% of the CMV consensus genome (percentage of identical matches: 90-100%). In Amiens, the analysis detected 9.79% of the HSV-1 consensus genome (percentage of identical matches: 91.66-97.5%) in two qPCR-confirmed individuals and 2.24% of the HHV-6A consensus genome (percentage of identical matches: 97.82%) in one qPCR-confirmed individual. Additionally, qPCR detection of albumin ranged from 36.6% in Amiens to 52.6% in Sevastopol; moreover, metagenomics identified two HSV-1 and one HHV-6 sequences in Amiens and two HSV-1, two HSV-2, one EBV and one VZV sequences in Sevastopol (Table 1, Figure 1).

**Table 1.** Detection of herpes aDNA in dental pulp samples collected from 16 individuals in an 17^th^ century burial site in Amiens, France, and from 36 individuals in a 19^th^ Crimean War burial site in Sevastopol, Crimea, and controls.

|  | Amiens<br>Teeth, n= 60 | Sevastopol<br>Teeth, n=112 |
| --- | --- | --- |
| Phage T4 | 60 (100%) | 111 (99.1%) |
| Albumin | 22 (36.6%) | 59 (52.6%) |
| HSV-1 | 3 (2) (5%) (2□) | 5 (4.5%) (2□) |
| HSV-2 | 0 (0%) | 2 (1.8%) |
| VZV | 0 (0%) | 1 (0.9%) |
| EBV | 0 (0%) | 1 (0.9%) |
| CMV | 0 (0%) | 0 (0%) |
| HHV-6 | 1 (1.7%) | 0 (0%) |
| HHV-7 | 0 (0%) | 0 (0%) |
| HHV-8 | 0 (0%) | 0 (0%) |

(2): 2 teeth belonging to the same individual, n: number of samples
□ Positive teeth with a negative albumin qPCR.

**Figure 1:**
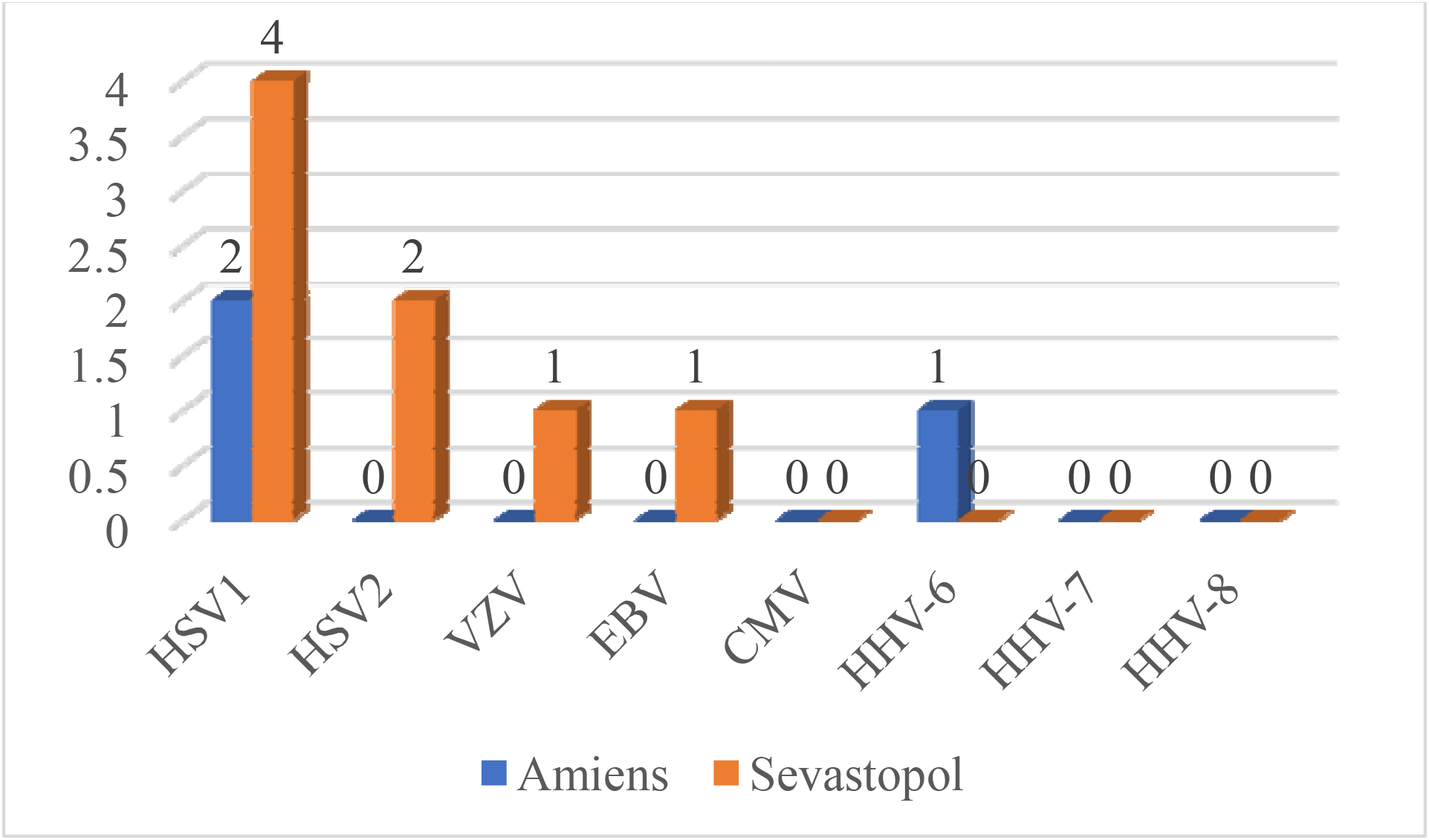
Detection of ancient Herpesviridae DNA in 17^th^ century individuals in Amiens (France) (blue bars) and in 19^th^ century individuals in Sevastopol (orange bars).

Furthermore, fluorescent *in situ* hybridization (FISH) targeting the HHV-6 UL67 gene in an individual in Amiens was performed as previously described, with some modifications (Atieh et al., 2013; Loukil, Kirtania, Bedotto, & Drancourt, 2018) (Appendix 3). The FISH probes V6-555: 5’-CGCTAGGTTGAGGATGATCGA-3’, V6-488: 5’-ATTCCTTCGGGTGTGACGTCTGGTG-3’ and V6-647: 5 ’-CGCTCTGGATAATTTGGCTTTG-3’, labeled with Alexa-555, Alexa-488 and Alexa-647 fluorochrome, respectively (Eurogentec, Angers, France), were applied, and an HHV-6 qPCR-negative sample from Amiens served as a negative control. FISH revealed fluorescent spots in the qPCR-confirmed sample but not in the negative control (Figure 2).

**Figure 2:**
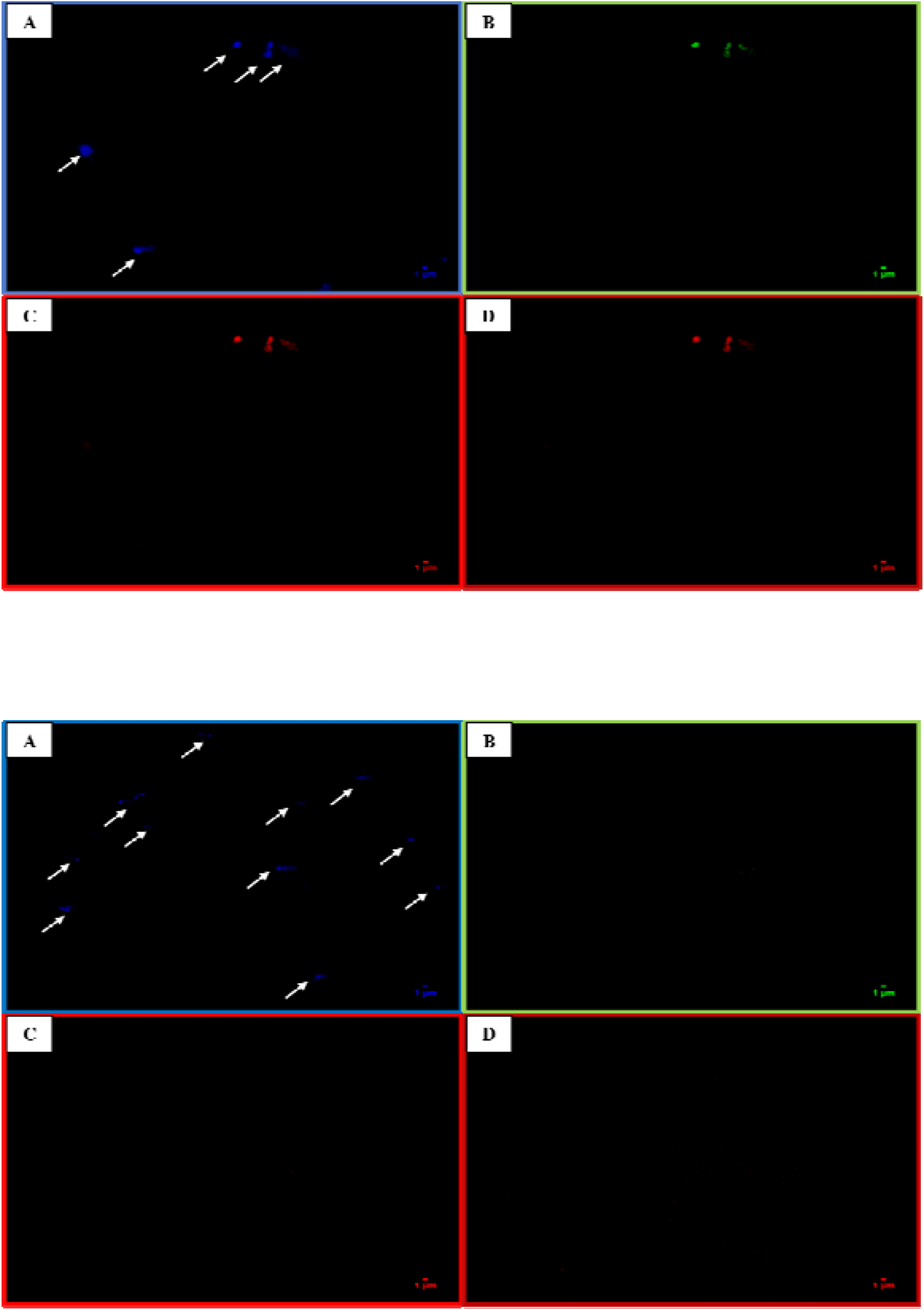
Panel A: FISH observation of HHV-6 aDNA in a dental pulp sample collected from an 17^th^ century individual in Amiens, France. The DAPI filter was used to visualize all the cells in the microscope field in blue (indicated with arrows, A); the FITC filter was used to visualize the probe V6-488 in green (B); the RHOD filter was used to visualize the probe V6-555 in red (C); and the CY5 filter was used to visualize the probe V6-647 in far-red. Panel B: FISH observation of the negative control using the same experimental conditions as above.

This pilot study offered the proof-of-concept that adapting the NovaSeq instrument and protocol for modern DNA metagenomics allowed detection of aDNA within a span of four days. Additionally, the NovaSeq instrument, having a weekly capacity of thousands of sequences, makes this instrument and packaged protocols an attractive first-line technique for pathogen detection in ancient specimens, which should be confirmed by deep sequencing of the entire pathogen genome after aDNA repair and paleoproteomics (Barbieri *et al*., 2017). Additionally, ancient dental pulp was a suitable material for such experiments, as it was previously used to identify hepatitis C virus in modern populations (Siravenha *et al*., 2016).

## ACKNOWLEDGMENTS

The authors acknowledge the personnel of the Sequencing Platform of IHU Méditerranée Infection. This work was supported by the French Government under the Investissements d’avenir (Investments for the Future) program managed by the Agence Nationale de la Recherche (ANR, fr: National Agency for Research), (reference: Méditerranée Infection 10-IAHU-03).

## Appendix 1.

### NovaSeq sequencing

In brief, genomic DNA was sequenced with the paired-end strategy and was barcoded for mixing with other genomic samples prepared with a Nextera XT DNA Sample Prep Kit (Illumina Inc.). To prepare the paired-end library, dilution was performed to obtain 1 ng of each genome as input. After tagmentation, limited 18-cycle PCR amplification completed the addition of the tag adapters and introduced dual-index barcodes. After purification with AMPure XP Beads (Beckman Coulter Inc., Fullerton, CA, USA), libraries were manually normalized. Library size was determined by a 2100 Bioanalyzer using a High-Sensitivity DNA Kit (Agilent, Waldbronn, Germany), and the concentration was determined by a Qubit dsDNA HS Assay Kit (Fisher Scientific, Carlsbad, USA). Library pooling (3 nM) and denaturation followed protocol A (standard loading) of the “NovaSeq 6000 System Denature and Dilute Libraries Guide” (document #1000000106351 v03). Then, a 15-hour sequencing run was performed with a NovaSeq 6000 SP Reagent Kit (100 cycles) v1.5 (Illumina, Inc.).

## Appendix 2.

### qPCR

Briefly, qPCR amplification of the albumin gene, phage T4, and the conserved regions of HHVs was performed in a 20-μL reaction volume including 15 μL of the mixture and 5 μL of extracted DNA or a 25-μL volume of the Argene kit reagent (bioMérieux, Marcy-l’Etoile, France) containing 15 μL of the mixture and 10 μL of extracted DNA. Every reaction also included cultured dental pulp stem cells (CLS, Eppelheim, Germany) and a reaction mixture as negative controls, and no positive controls were used. Amplification reactions were performed in a LightCycler 480 II real-time PCR system (Roche, La Rochelle, France) or a CFX96™ Real-Time System (Bio–Rad, Roanne, France) using the following protocol: one cycle of 95 °C for 10 min followed by 45 cycles of 95 °C for 15 s and 60 °C for 30 s.

**Table S1.**
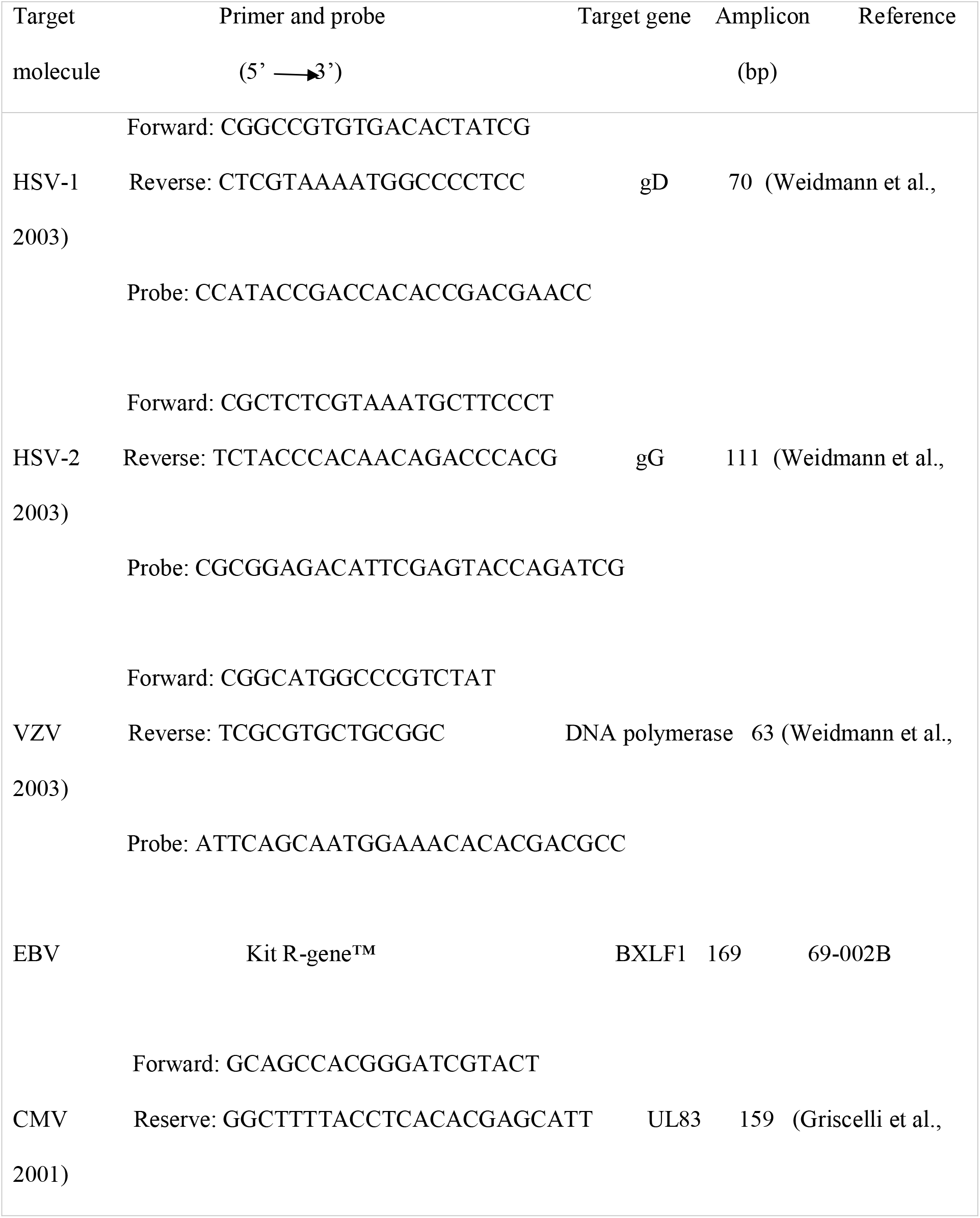
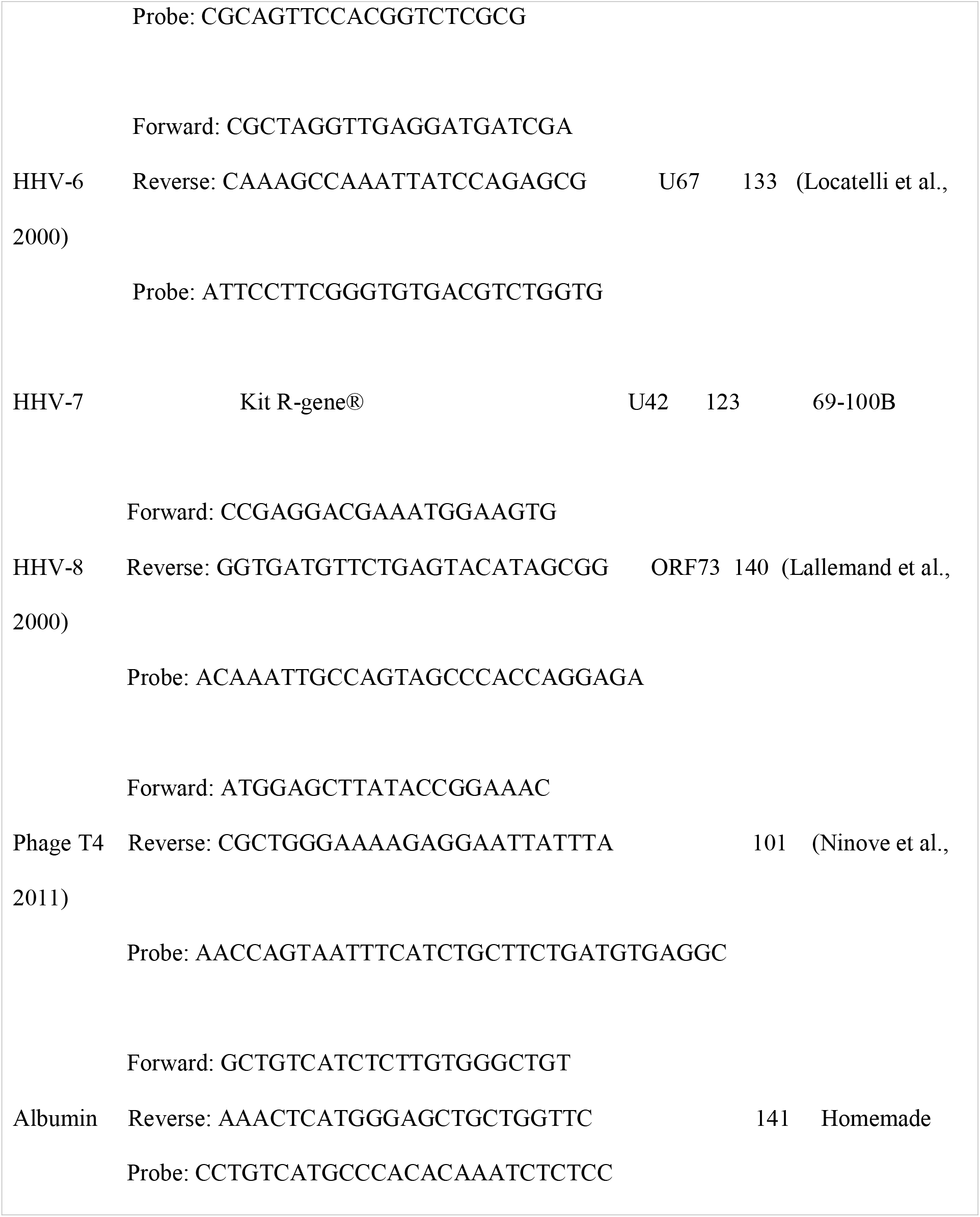
List of primers and probes used in this study for the detection of herpesvirus DNA in ancient dental pulp samples. All probes were labeled with FAM at the 3’ end and TAMRA at the 5’ end except phage T4 and albumin probes that were labeled by VIC at the end of 3’.

## Appendix 3.

### FISH

Briefly, 20 µL of rehydrated dental pulp was fixed with 4% paraformaldehyde on a glass slide and incubated for 5 min at 37 °C with 5 µg/mL proteinase K (Sigma–Aldrich, Saint-Quentin-Fancy, France). Then, 10 µL of probe solution (10 µmol/mL), hybridization buffer (a solution containing 0.1% Tween 20 and 0.1% Triton X-100; Euromedex, Souffelweyersheim, France), and distilled water were incubated for 10 min at 65 °C and then overnight at 37 °C. The slide was then immersed in a series of sodium saline citrate baths (4 ×, 2 ×, 1 ×, and 0.5 ×) for 5 min in each bath at room temperature (22 to 27 °C). Finally, a drop of Prolong Antifade mounting medium (Fisher Scientific, Illkirch, France) containing 4’,6’-diamidino-2-phenylindole was placed on the slide prior to microscopy using a fluorescence microscope (100× magnification, Leica DMI 6000, Nanterre, France).

